# Viruses inhibit TIR gcADPR signaling to overcome bacterial defense

**DOI:** 10.1101/2022.05.03.490397

**Authors:** Azita Leavitt, Erez Yirmiya, Gil Amitai, Allen Lu, Jeremy Garb, Benjamin R. Morehouse, Samuel J. Hobbs, Philip J. Kranzusch, Rotem Sorek

**Author notes:** These authors contributed equally to this work (Azita Leavitt, Erez Yirmiya, Gil Amitai, Allen Lu).

## Abstract

The Toll/interleukin-1 receptor (TIR) domain is a key component of immune receptors that identify pathogen invasion in bacteria, plants, and animals. In the bacterial antiphage system Thoeris, as well as in plants, recognition of infection stimulates TIR domains to produce an immune signaling molecule whose molecular structure remained elusive. This molecule binds and activates the Thoeris immune effector, which then executes the immune function. We identified a large family of phage-encoded proteins, denoted here Thoeris anti-defense 1 (Tad1), that inhibit Thoeris immunity. We found that Tad1 proteins are “sponges” that bind and sequester the immune signaling molecule produced by TIR-domain proteins, thus decoupling phage sensing from immune effector activation and rendering Thoeris inactive. A high-resolution crystal structure of Tad1 bound to the signaling molecule revealed that its chemical structure is 1′–2′ glycocyclic ADPR (gcADPR), a unique molecule not previously described in other biological systems. Our results define the chemical structure of a central immune signaling molecule, and reveal a new mode of action by which pathogens can suppress host immunity.

## Main Text

TIR-domains serve as the signal transducing modules in immune receptors that recognize pathogen invasion in the immune systems of bacteria, plants, and animals^1–3^. Whereas TIR domains in animals mainly transfer the signal by protein-protein interactions^3^, in plants and bacteria these domains produce an immune signaling molecule, which has the same mass as cyclic ADP-ribose (cADPR), but whose molecular and chemical structure remains elusive^1,4,5^. The mechanism of action of TIR-mediated immune signaling was recently deciphered for a bacterial anti-phage immune system called Thoeris^1^. This system comprises two core proteins, one of which (named ThsB) has a TIR domain and serves as the sensor for phage infection. Recognition of phage triggers the ThsB TIR domain to produce the cADPR isomer molecule, and this molecule activates a second Thoeris protein, ThsA, which then depletes the cell of the essential molecule nicotinamide adenine dinucleotide (NAD^+^) and leads to premature cell death to abort the infection^1,6^. Intriguingly, activation of plant TIRs by pathogens also leads to cell suicide that prevents pathogen propagation, and it was hypothesized that the elusive cADPR isomer produced by the TIRs is involved in mediating this plant immune response^4,5,7^.

### Identification of phage genes that inhibit Thoeris

We have isolated and analyzed a group of closely-related phages that infect *Bacillus subtilis*. This group included eight phages similar to phage SBSphiJ, a *Myoviridae* phage with a ∼150 kb-long genome^8^. Despite high sequence similarity between these phages, each phage had a few genes that were not found in the genomes of other phages in the group. (Figure 1A; Table S1). The Thoeris defense system protected against all phages from the SBSphiJ group except for phage SBSphiJ7 (Figure 1B). We therefore hypothesized that one or more genes that are unique to SBSphiJ7 allow this phage to escape or inhibit the activity of Thoeris.

**Figure 1.**
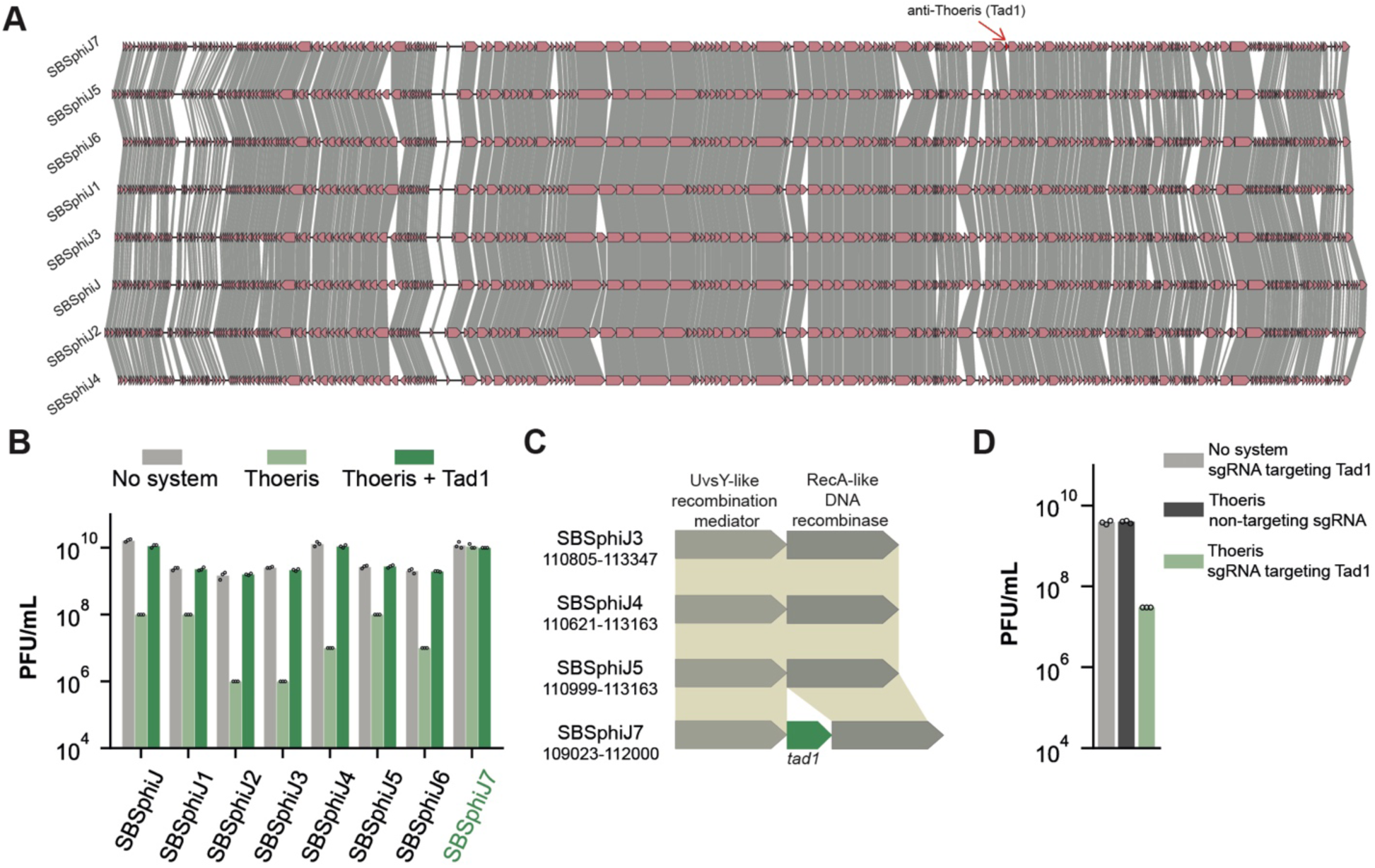
Tad1 inhibits Thoeris defense. (A) Genome comparison of eight phages from the SBSphiJ group. Amino acid sequence similarity between the ORFs is marked by grey shading. Genome similarity was visualized using clinker^10^. (B) Differential defense of Thoeris against SBSphiJ phages, and anti-Thoeris activity of Tad1. Data represent plaque-forming units per mL (PFU/mL) of phages infecting control cells (“no system”), cells expressing the Thoeris system (“Thoeris”), and cells co-expressing the Thoeris system and the *tad1* gene from SBSphiJ7. All phages except for SBSphiJ7 lack *tad1*. Shown is the average of three replicates, with individual data points overlaid. (C) The *tad1* locus in SBSphiJ7, shown in the locus for three other phages, SBSphiJ3, SBSphiJ4 and SBSphiJ5, in which *tad1* is absent. The coordinates of the presented locus within the phage genome are indicated below the name of each phage. (D) Tad1 knockdown cancels anti-Thoeris activity. Data represent PFU/mL of SBSphiJ7 that infects cells expressing Thoeris and a dCas9 system targeting Tad1, as well as control cells. Shown is the average of three replicates, with individual data points overlaid.

To test this hypothesis, we cloned five genes that were unique to phage SBSphiJ7, under the control of an inducible promoter, into *B. subtilis* cells that also express the Thoeris system from *Bacillus cereus* MSX-D12 (Table S2). One of these genes, which we denote *tad1* (Thoeris anti-defense 1) robustly inhibited the activity of Thoeris, as phages that were normally blocked by Thoeris were able to infect Thoeris-expressing cells if these cells also expressed Tad1 (Figure 1B, 1C). Silencing the expression of Tad1 in SBSphiJ7 using dCas9^9^ caused SBSphiJ7 to be blocked by Thoeris, verifying that *tad1* is the gene responsible for the Thoeris-inhibiting phenotype of SBSphiJ7 (Figure 1D).

Tad1 is a small protein (142 amino acids) of unknown function, with no recognizable protein domains. A search based on sequence homology identified 799 Tad1 homologs in the integrated microbial genomes (IMG)^11^ and the metagenomic gut virus (MGV)^12^ databases (Tables S3–S4). Strikingly, all homologs of Tad1 reside either in phage genomes or in genomes of prophages integrated within bacterial genomes, indicating that this protein primarily executes a phage function. Tad1-encoding phages belonged to multiple phage families, including *Myoviridae, Podoviridae* and *Siphoviridae*, and infect at least 60 species of bacteria from a diverse set of taxonomic phyla including *Proteobacteria, Firmicutes* and *Cyanobacteria* (Figure 2A; Tables S3– S4).

**Figure 2.**
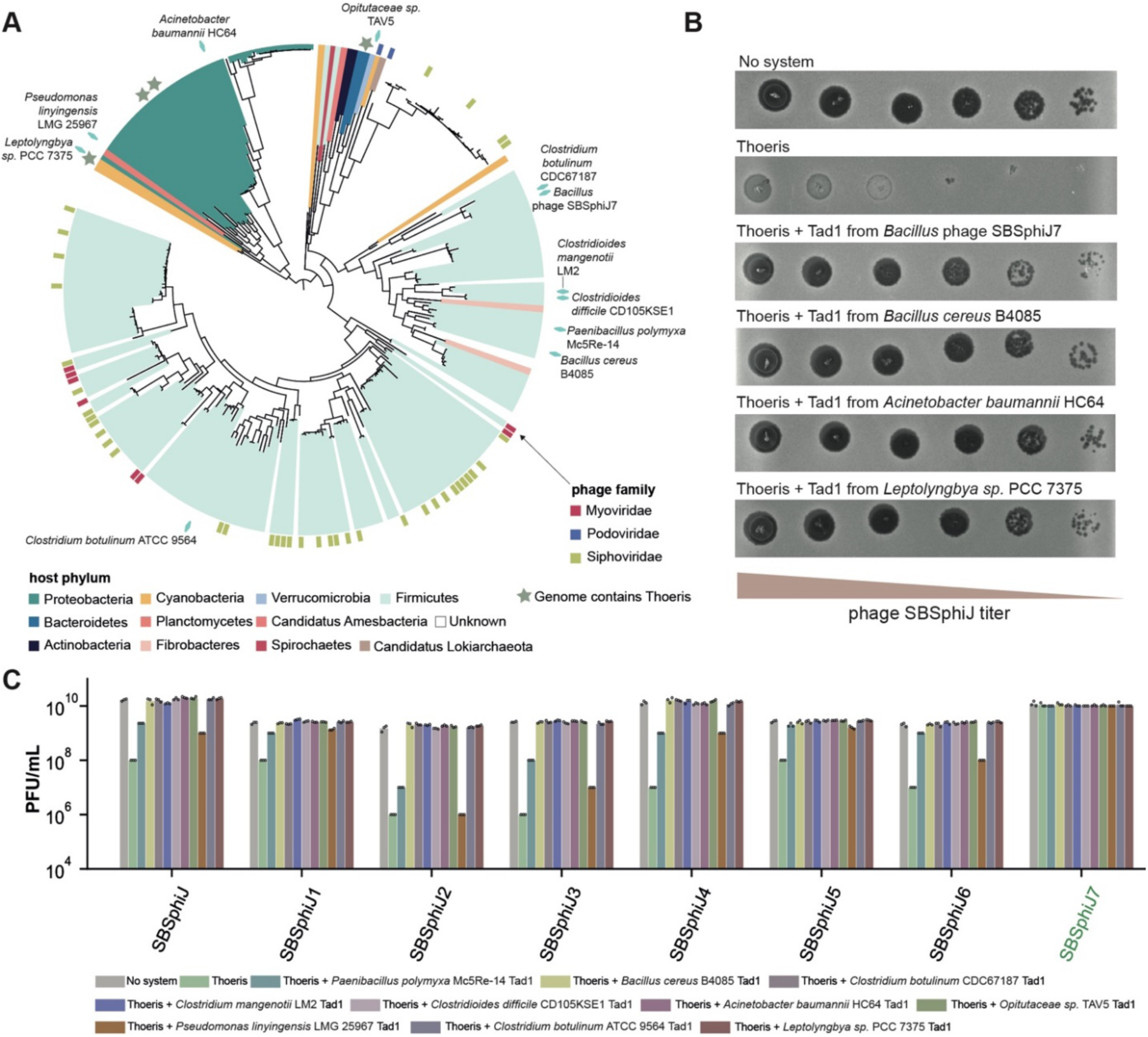
Tad1 proteins inhibit Thoeris defense. (A) Phylogenetic analysis of Tad1 homologs in phage and prophage genomes. The names of bacteria in which Tad1 homologs were found in prophages and tested experimentally are indicated on the tree as blue diamonds. Tree is based on 256 non-redundant sequences. (B) Tad1 homologs inhibit the Thoeris system in *B. subtilis*. Shown are tenfold serial dilution plaque assays with phage SBSphiJ. (C) Results of phage infection experiments with eight phages of the SBSphiJ family. Data represent PFU/mL of phages infecting control cells without Thoeris, cells expressing the Thoeris system, and cells co-expressing the Thoeris system and a Tad1 homolog. All phages except for SBSphiJ7 lack Tad1. Shown is the average of three replicates, with individual data points overlaid. The “Thoeris” and “no system” data presented here are the same as those presented in Figure 1B.

A phylogenetic analysis showed that Tad1 proteins can be divided into four major clades (Figure 2A). In some cases, phages encoding Tad1 proteins from the same phylogenetic clade belonged to different phage families, suggesting that *tad1* genes can be horizontally transferred between phages (Figure 2A). We selected ten Tad1 homologs that span the phylogenetic diversity of the Tad1 family, and cloned each of them into *B. subtilis* cells that express the Thoeris system (Figure 2A, S1). All ten Tad1 family proteins were able to inhibit Thoeris, including homologs derived from phages that infect distant organisms such as *Leptolyngbya sp*., *Opitutaceae sp*. and *Acinetobacter baumannii* (Figures 2A–C). Together, these results reveal a large family of proteins utilized by phages to inhibit the activity of the Thoeris bacterial defense system.

### Tad1 inhibits Thoeris by binding and sequestering TIR-derived signaling molecules

As most of the known anti-CRISPR proteins function by directly binding and inhibiting key proteins in the CRISPR-Cas complex^13–15^ we initially anticipated that Tad1 will bind one of the two Thoeris proteins, ThsA or ThsB. However, Tad1 did not co-immunoprecipitate with either ThsA or ThsB, implying that Tad1 inhibits Thoeris via a mechanism that does not require physical interaction with Thoeris proteins (Figure S2).

The hallmark of Thoeris defense is the TIR-domain ThsB protein which, upon sensing the infection, produces a signaling molecule that triggers the NADase activity of the Thoeris ThsA protein^1^. To test if this signaling molecule is produced in the presence of Tad1, we experimented with cells expressing a Thoeris system in which ThsA was mutated in its NADase active site such that only ThsB is active. We infected these cells with phage SBSphiJ that naturally lacks Tad1, and then lysed the cells and filtered the lysates to enrich for molecules smaller than 3 kDa (Figure 3A). As expected, purified ThsA protein incubated with these filtered lysates *in vitro* showed strong NADase activity, indicating that the TIR-domain ThsB protein produced the signaling molecule within the cell in response to SBSphiJ infection (Figure 3B). However, filtered lysates derived from cells in which Tad1 was co-expressed with ThsB failed to activate ThsA *in vitro*, suggesting that the signaling molecule was eliminated or inactivated in Thoeris-infected cells that co-express Tad1 (Figure 3B). Liquid chromatography followed by mass spectrometry (LC-MS) confirmed that the signaling molecule, having a mass identical to the mass of cADPR, was present in lysates of infected cells expressing the TIR-domain protein ThsB, but absent in lysates derived from infected cells where ThsB was co-expressed with Tad1 (Figure 3C). These results indicate that Tad1 does not inhibit the Thoeris effector protein ThsA, but rather inhibits Thoeris upstream of the ThsA protein.

**Figure 3.**
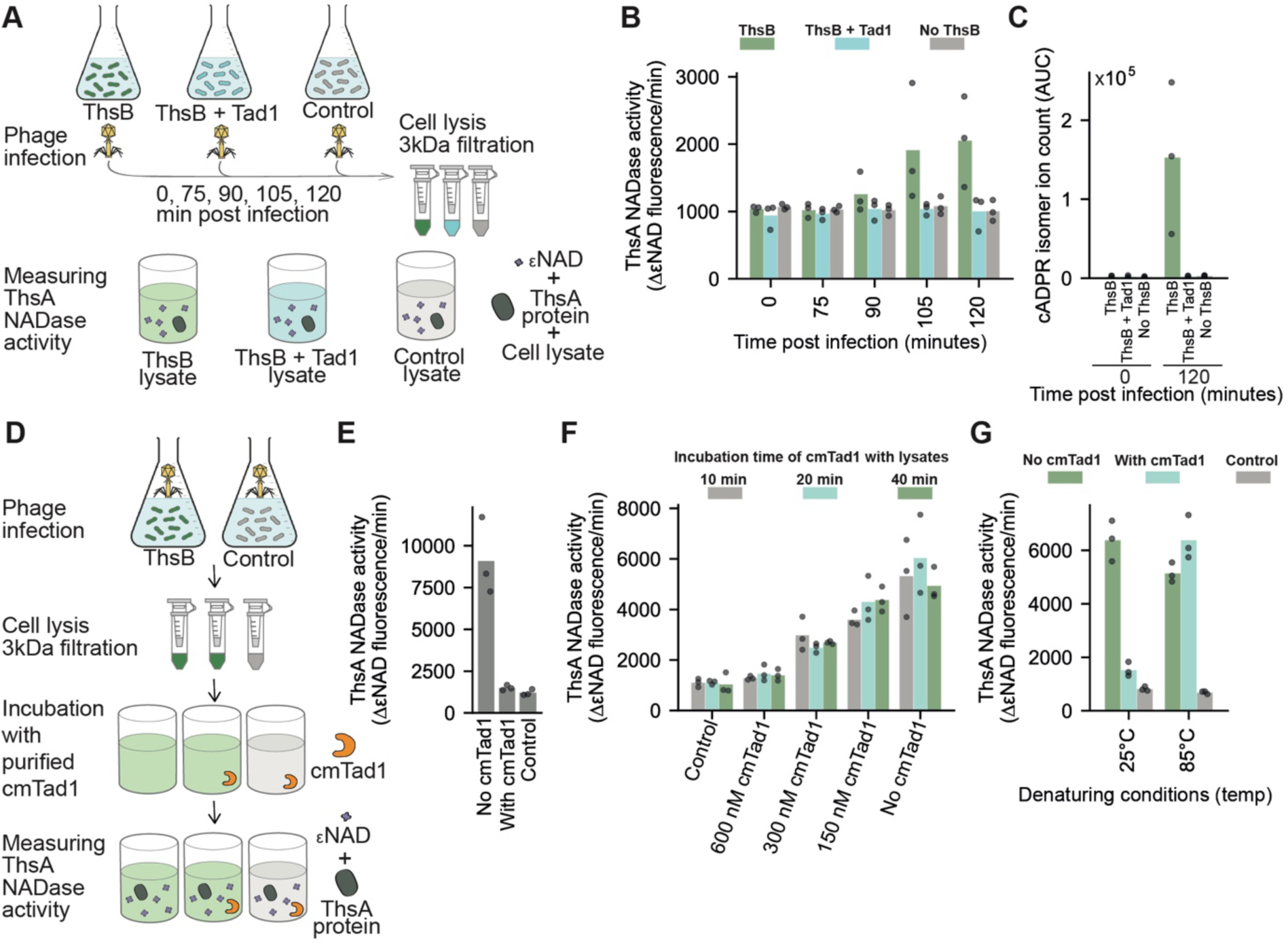
Tad1 cancels Thoeris-mediated defense by physically binding and sequestering the Thoeris-derived signaling molecule. (A) Schematic representation of the ThsB/Tad1 co-expression experiment. Cells expressing ThsB (native promoter), both ThsB (native promoter) and Tad1 (induced by 1 mM IPTG), or control cells that do not express ThsB, were infected with phage SBSphiJ at multiplicity of infection (MOI) of 5. NADase activity of purified ThsA incubated with filtered lysates was measured using a nicotinamide 1,N6-ethenoadenine dinucleotide (εNAD) cleavage fluorescence assay. (B) Activation of ThsA NADase activity by lysates from infected cells. Bars represent the mean of three experiments, with individual data points overlaid. (C) Co-expression of Tad1 with ThsB eliminates the signaling molecule normally produced by ThsB in infected cells. Lysates derived from infected cells were analyzed by LC-MS. Y axis represents the area under the curve (AUC) of cADPR isomer ions detected in the MS analysis. Time 0 represents uninfected cells. (D) Schematic representation of the Tad1/lysate incubation experiment. Cells overexpressing ThsB (0.1 mM IPTG) or control cells not expressing ThsB were infected with phage SBSphiJ at an MOI of 5 at 25°C. After 120 minutes the cells were lysed and lysates were filtered, and then incubated with purified cmTad1. ThsA was added to the lysates and NADase activity of ThsA was measured. (E) Purified Tad1 eliminates the signaling molecule from infected lysates. Filtered lysates were either pre-incubated with 600 nM purified cmTad1 for 10 minutes *in vitro* (“with cmTad1”) or with buffer (“no cmTad1”). (F) Tad1 is not an enzyme. Filtered lysates were incubated *in vitro* for 10, 20 or 40 minutes with purified cmTad1, or with a buffer, prior to exposure to ThsA. (G) Tad1 releases bound molecule when denatured. Shown is NADase activity of purified ThsA incubated with filtered lysates derived from infected cells overexpressing ThsB, that were additionally pre-incubated with purified cmTad1 *in vitro* for 10 minutes, followed by an additional incubation of 5 minutes at either 25°C or 85°C (denaturing conditions). “With cmTad1”, lysates pre-incubated with cmTad1. “No cmTad1”, lysates incubated with buffer instead of cmTad1. Control are lysates derived from infected cells that do not express ThsB.

We next experimented with a purified Tad1 homolog from a prophage integrated in *Clostridioides mangenotii* (cmTad1), which showed high stability when purified *in vitro*. To test if cmTad1 can directly eliminate the signaling molecule from the lysate, we collected filtered lysates from cells overexpressing ThsB that were infected by phage SBSphiJ and incubated these lysates with cmTad1 for 10 minutes (Figure 3D). Lysates incubated with purified cmTad1 completely lost the ability to induce the NADase activity of ThsA, demonstrating that Tad1 rapidly eliminates the signaling molecule from the filtered lysate rather than inhibiting its production (Figure 3E).

Some phages were previously shown to inhibit bacterial immune signaling, for example cyclic oligoadenylate signaling in type III CRISPR-Cas systems, by introducing enzymes that cleave the signaling molecules^16^. We therefore hypothesized that Tad1 is an enzyme that cleaves the immune signaling molecule of Thoeris. Under this hypothesis, one would expect Tad1 to deplete the signaling molecule in a time-dependent manner. However, counter to our hypothesis, incubation of sub-inhibitory concentrations of cmTad1 with the filtered lysate for prolonged time did not result in time-dependent increased depletion of the active molecule from the lysate (Figure 3F). These results implied that Tad1 is not an enzyme, but rather a “sponge” that binds and sequesters the signaling cADPR isomer molecule.

To further examine the hypothesis that Tad1 tightly binds the cADPR isomer signaling molecule, we tested whether cmTad1 denaturation could release the bound molecule into the buffer. For this, we first incubated cmTad1 with the signal-containing lysate for 10 minutes, and then denatured it by exposure to 85°C for 5 minutes. We found that following Tad1 denaturation, the buffer regained the capacity to activate ThsA *in vitro* (Figure 3G). These results demonstrate that Tad1 binds and sequesters, but does not degrade, the Thoeris signaling molecule, and that denaturation of Tad1 releases the bound molecule intact. Our results therefore show that Tad1 inhibits Thoeris defense by physically binding and sequestering the ThsB-derived signaling molecule, thus preventing the activation of the Thoeris immune effector and mitigating Thoeris-mediated defense.

It was previously shown that cADPR isomer molecules produced by plant TIR proteins can activate the Thoeris effector ThsA^1^. We therefore co-expressed cmTad1 with a TIR-domain protein from the plant *Brachypodium distachyon* (BdTIR), which was shown to constitutively produce the cADPR isomer molecule when expressed in *E. coli*^4^. cmTad1 purified from BdTIR-expressing cells showed substantial shifts during size-exclusion chromatography and exhibited increased absorption at UV_260_ as compared to cmTad1 purified from control cells, suggesting that cmTad1 binds the signaling molecule produced by the plant TIR (Figure S3 A,B). Indeed, supernatant collected from heat-denaturation of the molecule-bound cmTad1 was able to activate ThsA, verifying that the signaling molecule produced by the plant TIR, which was bound by cmTad1, is the same molecule utilized by the Thoeris system (Figure S3 C,D).

### Crystal structure of Tad1 reveals the immune signal 1′–2′ glycocyclic ADPR (gcADPR)

To define the molecular mechanism of Tad1 anti-Thoeris evasion, we next determined crystal structures of Tad1 from a *Clostridium botulinum* prophage (cbTad1) in the apo and ligand-bound states (Supplementary Table 5). The structure of Tad1 reveals a homodimeric complex with two protomers arranged head-to-tail (Figure 4A). Each protomer forms an N-terminal anti-parallel β-sheet (β1–β4) and two long C-terminal helices (α1 and α2) that create a wedge-shaped architecture and allow Tad1_a_ and Tad1_b_ to tightly interlock into a compact assembly (Figure 4A,B). The N-terminal β-sheets join through a β4_a_–β4_b_ hydrophilic seam to form the front face of the Tad1 assembly, while the C-terminal helices align to create a four-helix bundle that seals the back face (Figure 4A,B). The tightly locked assembly creates two recessed ligand binding pockets at the top and bottom ends of the Tad1 complex that are each surrounded by four highly-conserved loops within β2–β3, β4–α1, and the C-terminal tail of Tad1_a_ along with α1–α2 donated by the partner protomer Tad1_b_ (Figure 4B, S1).

**Figure 4.**
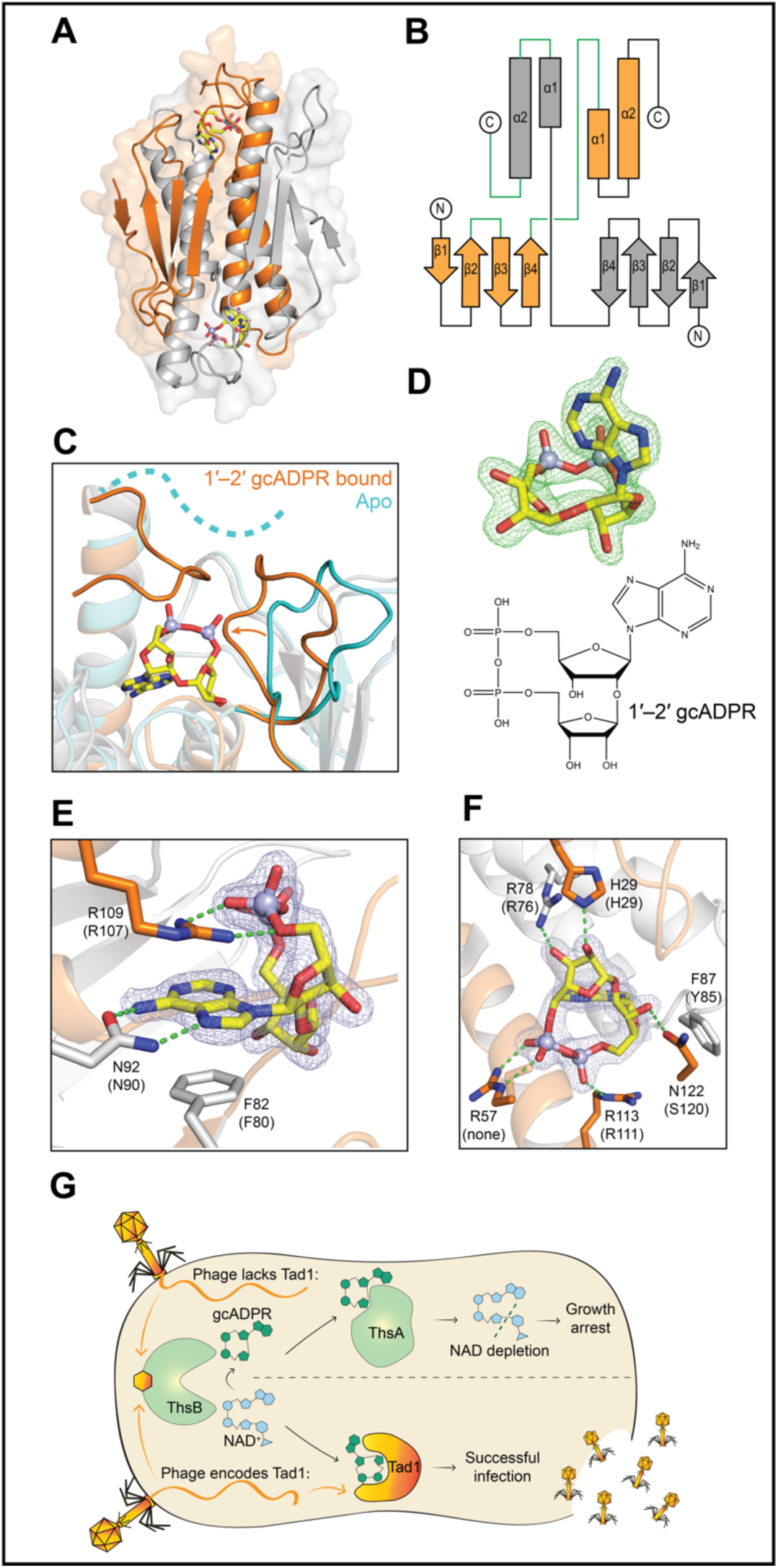
Structure of Tad1 and identification of 1’–2’ gcADPR. (A) Overview of the cbTad1 crystal structure in front view bound to 1’–2’ gcADPR (yellow). Tad1 forms a homodimer with two ligand binding sites, with one monomer shown in orange and the other in grey. (B) Topology map of Tad1 from a top view perspective. Loops that form one binding site are highlighted in green. (C) Comparison of the ligand binding site of cbTad1 in the *apo* state (cyan) and in complex with the Thoeris signal (orange and grey). Upon ligand binding, cbTad1 loop β4–α1 shifts ∼3.7 Å ⍰(measured by A56 amine movement) and the C-terminal tail becomes structured to enclose around the molecule. (D) Polder omit map of the Tad1 ligand-binding site contoured at 6σ reveals the chemical structure of the ligand as 1’–2’ gcADPR. (E) Detailed view of cbTad1 residues interacting with the adenine base of 1’–2’ gcADPR or (F) with the ribose moieties and phosphates. Conserved residues in cmTad1 are labeled separately in parentheses. Separate monomer chains are shown in either orange or grey. Green dashed lines denote hydrogen bonding interactions. Light blue mesh denotes the 1’–2’ gcADPR *2F*_*o*_ – *F*_*c*_ electron density contoured at 1.8*σ*. (G) A model of the mechanism of Tad1.

Upon ligand binding, the Tad1 complex undergoes a 3° rotation to close and envelope the TIR-derived signaling molecule. Tad1 loop β4–α1 moves >3.5 Å and the C-terminal residues 116–122 form an ordered lid that together seal the ligand-binding site (Figure 4C). Exceptionally clear density within the 1.9 Å ligand-bound cbTad1 complex allowed for unambiguous assignment of each atomic position within the TIR-derived signaling molecule, revealing the compound 1′–2′ glycosyl cyclic adenosine diphosphate ribose (1′–2′ gcADPR) (Figure 4D). In contrast to the canonical cADPR, in which cyclization occurs through the ribose and the N1 position in the adenine nucleobase, our data demonstrate that the signaling molecule produced by bacterial and plant TIR proteins is unexpectedly cyclized through a unique ribose–ribose glycosidic bond. The Tad1 ligand-binding pocket intimately embraces 1′–2′ gcADPR with 11 residues forming base- and linkage-specific contacts (Figure 4E,F). The 1′–2′ gcADPR adenine base is stacked between cbTad1 F82 and R109, with N92 making sequence-specific contacts to the Hoogsteen edge (Figure 4E). The diphosphate backbone is bound by three cbTad1 sidechains R57, R109, and R113, and additional peptide-backbone contacts from R57 and G120. Finally, cbTad1 F87 buttresses the adenosine ribose and residues H29, G55(NH), R78, and N122 coordinate each free OH in the ribose–ribose linkage, explaining the intimate specificity of Tad1 for the unique linkage in 1′–2′ gcADPR (Figure 4F). Complete enclosure within the ligand-binding pocket explains how Tad1 efficiently sequesters the TIR-derived signal to inactivate Thoeris defense.

Together, our data reveal a complete mechanism of viral subversion from TIR-derived signaling through signal molecule sequestration (Figure 4G). We envision that similar mechanisms may be employed by plant pathogens to inhibit TIR-mediated plant immunity, and that signal sequestration could be a general mechanism utilized by viruses to evade immune responses that rely on signaling molecules. Our results establish 1′–2′ gcADPR as a signaling molecule produced by bacterial and plant immune TIR proteins. This molecule is unique, having a chemical structure which to our knowledge was not previously described in any biological system. As in other immune pathways that employ signaling molecules^17–19^, we envision that future studies will reveal immune systems with TIR-derived signals that deviate from the archetypal 1′–2′ gcADPR molecule. Finally, as recent studies exposed numerous antiviral mechanisms that are conserved from bacteria to humans^19–22^ it would be intriguing to examine whether the human immune system also involves gcADPR signaling.

## Methods

### Phage strains, isolation, cultivation and sequencing

Phage SBSphiJ was isolated in a previous study^8^. Other phages used in this study were isolated from soil samples on *B. subtilis* BEST7003 culture as described in Doron *et al*^8^. For this, soil samples were added to a log phase *B. subtilis* BEST7003 culture and incubated overnight to enrich for *B. subtilis* phages. The enriched samples were centrifuged and filtered through 0.45 µm filters, and the filtered supernatant was used to perform double layer plaque assays as described in Kropinski *et al*.^23^. Single plaques that appeared after overnight incubation were picked, re-isolated three times, and amplified as described below.

Phages were propagated by picking a single phage plaque into a liquid culture of *B. subtilis* BEST7003 grown at 37°C to OD_600_ of 0.3 in magnesium manganese broth (MMB) (LB + 0.1 mM MnCl_2_ + 5 mM MgCl_2_) until culture collapse. The culture was then centrifuged for 10 minutes at 3,200 × *g* and the supernatant was filtered through a 0.2 µm filter to remove remaining bacteria and bacterial debris.

High titer phage lysates (>10^7^ pfu/mL) were used for DNA extraction. 500 µL of the phage lysate was treated with DNase-I (Merck cat #11284932001) added to a final concentration of 20 mg mL^−1^ and incubated at 37°C for 1 h to remove bacterial DNA. DNA was extracted using the QIAGEN DNeasy blood and tissue kit (cat #69504) starting from the Proteinase-K treatment step to lyse the phages. Libraries were prepared for Illumina sequencing using a modified Nextera protocol as previously described^24^.

Following Illumina sequencing, adapter sequences were removed from the reads using Cutadapt version 2.8^25^ with the option -q 5. The trimmed reads from each phage genome were assembled into scaffolds using SPAdes genome assembler version 3.14.0^26^, using the –careful flag. Each assembled genome was analyzed with Prodigal version 2.6.3^27^ (default parameters) to predict ORFs.

### Plaque assays

Phage titer was determined using the small drop plaque assay method^28^. 400 μL of the bacterial culture was mixed with 30 mL melted MMB 0.5% agar, poured on 10 cm square plates, and let to dry for 1 h at room temperature. In cases of bacteria expressing anti-defense candidates, 1 mM IPTG was added to the medium. In cases of bacteria expressing dCas9-gRNA constructs, 0.002% xylose was added to the medium. 10-fold serial dilutions in MMB were performed for each of the tested phages and 10 µL drops were put on the bacterial layer. After the drops had dried up, the plates were inverted and incubated at room temperature overnight. Plaque forming units (PFUs) were determined by counting the derived plaques after overnight incubation and lysate titer was determined by calculating PFUs per mL. When no individual plaques could be identified, a faint lysis zone across the drop area was considered to be 10 plaques. Efficiency of plating (EOP) was measured by comparing plaque assay results on control bacteria and bacteria containing the defense system and/or a candidate anti-defense gene.

### Prediction and cloning of the candidate anti-Thoeris genes

Predicted protein sequences from all phage genomes were clustered into groups of homologs using the cluster module in MMSeqs2 release 12-113e3 ^29^, with the parameters -e 10, -c 0.8, -s 8, --min-seq-id 0.3 and the flag –single-step-clustering. Anti-Thoeris candidates were defined as clusters that included a single member derived from phage SBSphiJ7. The DNA of each anti-Thoeris candidate was amplified from the genome of phage SBSphiJ7 using KAPA HiFi HotStart ReadyMix (Roche cat # KK2601) with primers as indicated in Table S2. Homologs of *tad1* were synthesized by Genscript Corp. The anti-Thoeris candidates were cloned into the pSG-*thrC*-Phspank vector (Supplementary File S1) and transformed to 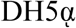 competent cells. The cloned vector was subsequently transformed into *B. subtilis* BEST7003 cells containing the Thoeris defense system^8^, resulting in cultures expressing both Thoeris and an anti-Thoeris gene candidate. As a negative control, a transformant with an identical plasmid containing GFP instead of the anti-Thoeris gene was used. Transformation to *B. subtilis* was performed using MC medium as previously described^8^. Whole-genome sequencing was then applied to all transformed *B. subtilis* strains, and Breseq analysis^30^ was used to verify the integrity of the inserts and lack of mutations.

### Construction of dCas9 and gRNA cassette for integration to *Bacillus Subtilis* thrC site

A shuttle vector was constructed for the integration of dCas9 from *Streptococcus pyogenes* into the thrC gene of *B. subtils*. The dCas9 gene, including its xyl promoter and xylR was amplified from pJMP1 (Addgene plasmid #79873) using primers “CTTCTAGGATCAAATCGATATCTCTGCAGTCGCG” and “AAAAATCCTTTTCTTTCTTATCTTGTGATTGGTGTATCATTTCGTTTTTCTTTTGTGC”.

The gRNA with a constitutive *B. subtilis* promoter was amplified from pJMP3 (Addgene plasmid #79875) using primers “AGATATCGATTTGATCCTAGAAGCTTATCGAATTCCTTATTAACGT” and “CGAATTCGACTCTCTAGCTCTACCATCGGCGCTACGG”. The two fragments were cloned into a backbone amplified from pSG_thrC_Phspank_sfGFP^31^ using primers “CAAGATAAGAAAGAAAAGGATTTTTCGCTACG” and “AGCTAGAGAGTCGAATTCGGATCCG”, resulting in a plasmid named pJG_thrC_dCAS9_gRNA (Supplementary File S2).

To insert new spacers, two fragments were amplified from pJG_thrC_dCAS9_gRNA and the new spacer was introduced in the overlap of primers designed for NEBuilder HiFi DNA Assembly (NEB, cat # E2621). For the gRNA used to target *tad1*, the first fragment was amplified using primers “TTCAACAAACGAAAATTGGATAAAGTGGGAT” and “GAACCACTACGAAATGATGGTTTTAGAGCTAGAAATAGCAAGTTAAAATAAGGCT”, and the second fragment was amplified using primers “CCAATTTTCGTTTGTTGAACTAATGGGTG” and “CATCATTTCGTAGTGGTTCCACATTTATTGTACAACACGAGCCCATTT”. The resulting assembled construct had the gRNA sequence ‘GGAACCACTACGAAATGAT’.

The gRNA sequence ‘CTATGATTGATTTTTTTAGC’ was used as a control. It was constructed as mentioned above, with primers “TTCAACAAACGAAAATTGGATAAAGTGGGAT” and “TGTCTATGATTGATTTTTTTAGCGTTTTAGAGCTAGAAATAGCAAGTTAAAATAAGGCT”, and “CCAATTTTCGTTTGTTGAACTAATGGGTG” and “GCTAAAAAAATCAATCATAGACATTTATTGTACAACACGAGCCCATTT”. The resulting vectors containing the dCas9-gRNA sequences were cloned to a *B. subtilis* strain that contained the Thoeris defense system, as well as to a control strain that lacks Thoeris. Shuttle vectors were propagated in *E. coli* DH5a with 100 μg/mL ampicillin selection. Plasmids were isolated from *E. coli* DH5a prior to transformation into the appropriate *B. subtilis* BEST7003 strains.

### Identification of anti-Thoeris homologs and phylogenetic reconstruction

Tad1 homologs were searched for in the metagenomic gut virus database (MGV)^12^ and in ∼38000 prokaryotic genomes downloaded from the IMG database^11^ in October 2017. The search was preformed using the “search” option of MMseqs release 12-113e3 with default parameters. The unique (non-redundant) sequences were used for multiple alignment with MAFFT version 7.402 ^32^ using default parameters. The phylogenetic tree was constructed using IQ-TREE version 1.6.5 ^33^ with the -m LG parameter. The online tool iTOL24 (v.5) ^34^ was used for tree visualization. Phage family annotations were based on the prediction in the MGV database. The host phyla annotations were either based on the prediction in the MGV database, or the IMG taxonomy of the bacteria in which the prophage was found.

### Pulldown attempts of ThsA and ThsB with tagged Tad1

*B. subtilis* phage SBSphiJ7 *tad1* gene was cloned into the pBbA6c plasmid containing a C-terminal 8xHis-tag and a lac promoter^35^ (Supplementary File S3). The *Bacillus cereus* MSX-D12 Thoeris *thsA* and *thsB* genes were cloned as an operon under the arabinose-inducible promoter on pBAD (Supplementary File S4). Both plasmids were co-transformed into an *E. coli* MG1655 strain.

Overnight cultures containing the pBAD-*thsA*-*thsB* and pBbA6c-*tad1*-His constructs were diluted 1:100 in 100 mL MMB and grown at 25°C, 200 rpm shaking. 1 mM IPTG and 0.2% arabinose were added at an OD_600_ of 0.3 and harvested after 3 h by centrifugation at 4000 RPM for 20 min at 4°C. Pellets were stored at −80°C and then lysed with 250 μL 50 mM Na-Phosphate buffer pH 8.0, 0.1 M NaCl and 1 mg mL^−1^ lysozyme (Sigma cat # L6876). Pellets resuspended with lysozyme were shaken for 10 min at 25°C and then 750 μL of NTA-washing buffer (Na-Phosphate buffer 20 mM pH 7.4, 0.5 M NaCl, 20 mM imidazole, 0.05% Tween 20) was added. The samples were transferred to a FastPrep Lysing Matrix B in a 2 mL tube (MP Biomedicals cat # 116911100) and lysed using FastPrep bead beater for 2 × 40 s at 6 m s^−1^. Tubes were then centrifuged at 4°C for 10 min at 15,000 × *g*.

The supernatant was then applied to Ni-NTA-magnetic beads (Qiagen 36113) and washed twice with 0.5 mL NTA-washing buffer, followed by elution with NTA-washing buffer containing 500 mM imidazole. Protein samples containing Tad1-His co-expressed with ThsA and ThsB were compared to cells expressing either Tad1-His or ThsA/ThsB by running the purified samples on a Bolt™ 4 to 12%, Bis-Tris protein gel (Thermofisher NW04122BOX).

### Preparation of filtered cell lysates

For the *in vivo* experiments, we used *B. subtilis* BEST7003 co-expressing a Thoeris system in which ThsA was mutated in its NADase active site (ThsA_N112A_+ThsB)^1^ under its native promoter, and Tad1 under the Physpank promoter. Controls included cells expressing only the mutated Thoeris system, as well as cells lacking both the Thoeris system and Tad1. These cultures were grown overnight and then diluted 1:100 in 350 mL MMB supplemented with 1 mM IPTG and grown at 37°C, 200 rpm shaking for 90 min. Each culture was then incubated and shaken at 25°C 200 rpm until reaching an OD_600_ of 0.3. At this point, a sample of 50 mL was taken as the uninfected (time 0 min) sample, and SBSphiJ phage was added to the remaining 300 mL culture at an MOI of 5. Flasks were incubated at 25°C with shaking (200 rpm), for the duration of the experiment. 50 mL samples were collected at time points 75, 90, 105, and 120 min post-infection. Immediately upon sample removal (including time point 0 min), the 50 mL sample tubes were placed on ice and centrifuged at 4°C for 10 min to pellet the cells. The supernatant was discarded and the pellet was flash frozen and stored at −80°C.

For the *in vitro* experiment with purified cmTad1, *Bacillus subtilis* BEST7003 cultures overexpressing the Thoeris ThsB protein under the Physpank promoter, together with control cultures, were diluted 1:100 in 200 mL MMB supplemented with 0.1 mM IPTG and grown at 37°C, with shaking at 200 rpm. After 90 min, temperature was lowered to 25°C and shaken (200 rpm) until reaching an OD_600_ of 0.3. Then, SBSphiJ phage was added to the culture at an MOI of 5. Flasks were incubated at 25°C with shaking (200 rpm). 120 min post infection the culture was collected into 50 mL tubes, centrifuged for 10 min at 4°C, supernatant discarded and pellet was stored at −80°C.

To extract the cell metabolites from frozen pellets, 600 µL of 100 mM phosphate buffer, pH 8.0, supplemented with 4 mg mL^−1^ lysozyme (Sigma cat # L6876) was added to each pellet. Tubes were then incubated for 10 min at 25°C, and returned to ice. The samples were transferred to a FastPrep Lysing Matrix B in a 2 mL tube (MP Biomedicals cat # 116911100) and lysed using FastPrep bead beater for 2 × 40 s at 6 m s^−1^. Tubes were then centrifuged at 4°C for 10 min at 15,000 × *g*. Supernatant was transferred to Amicon Ultra-0.5 Centrifugal Filter Unit 3 kDa (Merck Millipore cat # UFC500396) and centrifuged for 45 min at 4°C, 12,000 × *g*. Filtered lysates were taken either for LC-MS analysis or for in vitro NADase activity assays.

### LC–MC monitoring of the Thoeris cADPR isomer

Sample analysis was carried out by MS-Omics as follows. Samples were diluted 1:3 in 10% ultra-pure water and 90% acetonitrile containing 10 mM ammonium acetate at pH 9 then filtered through a Costar Spin-X centrifuge tube filter 0.22 μm Nylon membrane. The analysis was carried out using a UPLC system (Vanquish, Thermo Fisher Scientific) coupled with a high-resolution quadrupole-orbitrap mass spectrometer (Q Exactive™ HF Hybrid Quadrupole-Orbitrap, Thermo Fisher Scientific). The standard cADPR peak was identified using a synthetic standard (cADPR: Sigma-Aldrich, C7344) run. The UPLC was performed using an Infinity Lab PoroShell 120 hydrophilic interaction chromatography (HILIC-Z PEEK) lined column with dimensions of 2.1 × 150 mm and a particle size of 2.7 μm (Agilent Technologies). The composition of mobile phase A was 10 mM ammonium acetate at pH 9 in 90% acetonitrile LC–MC grade (VWR Chemicals) and 10% ultra-pure water from Direct-Q 3 UV Water Purification System with LC-Pak Polisher (Merck KGaA) and mobile phase B was 10 mM ammonium acetate at pH 9 in ultra-pure water with 15 μM medronic acid (InfinityLab Deactivator additive, Agilent Technologies). The flow rate was kept at 250 μL mL^−1^ consisting of a 2 min hold at 10% B, increased to 40% B at 14 min, held till 15 min, decreased to 10% B at 16 min and held for 8 min. The column temperature was set at 30°C and an injection volume of 5 μL. Analysis was performed in positive ionization mode from m/z 200 to 1000 at a mass resolution of 120,000 (at m/z 200). An electrospray ionization interface was used as ionization source. Peak areas were extracted using TraceFinder 4.1 (ThermoFisher Scientific) with an accepted deviation of 5 ppm. Fragmentation was done through a higher-energy collisional dissociation cell using a normalized collision energy of 20, 40 and 60 eV where the spectrum is the sum of each collision energy.

### CmTad1 protein cloning, expression, and purification

*Clostridioides mangenotii* Tad1 (cmTad1) was used for the *in-vitro* experiments because its higher stability enabled long term storage in −80°C. The *cmTad1* gene was cloned into a the expression vector pET28-bdSumo using the Restriction-Free (RF) method^36^ (Supplementary File S5). pET28-bdSumo was constructed by transferring the His14-bdSUMO cassette from the K151 expression vector generously obtained from Prof. Dirk Görlich from the Max-Planck-Institute, Göttingen, Germany^37^ into the expression vector pET28-TevH^38^.

CmTad1 was expressed in *E. coli* BL21(DE3) by induction with 200 μM IPTG at 15°C overnight. The culture was harvested and lysed by a cooled cell disrupter (Constant Systems) in a lysis buffer (50 mM Tris pH 8, 0.5 M NaCl, 1 mM DTT, 2 mM MgCl_2_, 250 mM Sucrose) containing 200 KU 100 mL^−1^ lysozyme, 20 μg mL^−1^ DNase, 1 mM phenylmethylsulfonyl fluoride (PMSF) and protease inhibitor cocktail (Millipore, 539134). After centrifugation, the supernatant of the lysate was incubated with 5 mL Nickel-beads (Adar Biotech, prewashed with lysis buffer) for 1 h at 4°C. After removing the supernatant by centrifugation, the beads were washed 3 × with 50 mL lysis buffer and once with lysis buffer containing 50 mM imidazole (1 mM TCEP replacing the DTT). CmTad1 was eluted from the beads by incubation with 20 mL cleavage buffer (50 mM Tris pH 8, 0.5 M NaCl, 1 mM DTT, 2 mM MgCl_2_, 250 mM Sucrose and 0.4 mg bdSumo protease) for 2 h at 25°C. The supernatant containing the cleaved cmTad1 was removed and an additional 20 mL cleavage buffer was added to the beads and left overnight at 4°C. The two elution samples were combined, concentrated, and applied to a size exclusion column (HiLoad 16/60 Superdex200 prep-grade, Cytiva) equilibrated with a SEC buffer (50 mM Tris pH 8, 50 mM NaCl, 1 mM DTT). Fractions containing pure cmTad1 were pooled and frozen at −80°C.

### NADase activity assay as a reporter for the cADPR isomer

NADase assay was performed by using the *Bacillus cereus* MSX-D12 ThsA enzyme as a reporter for the presence of the cyclic ADPR isomer^1^. The reporter enzyme ThsA was expressed and purified as described previously^1^. NADase reaction was performed in black 96-well half area plates (Corning cat # 3694) at 25°C in a 50 µl final reaction volume. 5 µL of 5 mM nicotinamide 1,N6-ethenoadenine dinucleotide (εNAD, Sigma, N2630) solution was added to each well sample immediately prior to measurements and mixed by pipetting. εNAD was used as a fluorogenic substrate to report the ThsA enzyme NADase activity by monitoring increase in fluorescence (excitation 300 nm / emission 410 nm) using a Tecan Infinite M200 plate reader at 25°C. Reaction rate was derived from the linear part of the initial reaction.

For the assessment of the *in vivo* activity of Tad1 in cells co-expressing both ThsB (native promoter) and Tad1, filtered lysates were mixed directly with ThsA, followed by the addition of εNAD. Controls included filtered lysates derived from cells expressing ThsB (without Tad1), and cells expressing neither ThsB nor Tad1. For the assessment of the *in vitro* activity of purified cmTad1, phage-infected cell filtered lysate (diluted 1:16–1:20) overexpressing ThsB (100μM IPTG) were incubated with purified cmTad1 (150–600 nM final concertation) at 25°C. Controls included filtered lysates incubated with diluted SEC buffer.

To examine the level of the cADPR isomer after cmTad1 heat-inactivation, cmTad1 (600nM) was incubated with phage-infected cell filtered lysate (diluted 1:16) for 10 min at 25°C followed by incubation at either 85°C or 25°C for an additional 5 min. Samples were then mixed with 2 µL of 2.5 μM ThsA and 5 µL of 5 mM εNAD and fluorescence was monitored as descried above.

### Tad1 crystallization and structural analysis

For crystallography experiments, *cmTad1* and *cbTad1* genes were cloned from synthetic DNA fragments (Integrated DNA Technologies) by Gibson assembly into a custom pET expression vector containing an N-terminal 6×His-SUMO2 tag as previously described^39^. Plasmids were transformed into BL21(DE3) RIL *E. coli* (Agilent) and colonies were grown on MDG agar plates (1.5% agar, 2 mM MgSO_4_, 0.5% glucose, 25 mM Na_2_HPO_4_, 25 mM KH_2_PO_4_, 50 mM NH_4_Cl, 5 mM Na_2_SO_4_, 0.25% aspartic acid, and 2–50 μM trace metals). Three colonies were picked into a 30 mL MDG starter culture and grown overnight at 37°C with 230 rpm shaking. 15 mL of overnight cultures were used to seed 1 L M9ZB expression cultures (2 mM MgSO_4_, 0.5% glycerol, 47.8 mM Na_2_HPO_4_, 22 mM KH_2_PO_4_, 18.7 mM NH_4_Cl, 85.6 mM NaCl, 1% Cas-Amino acids, 2– 50 μM trace metals, 100 μg mL^−1^ ampicillin, and 34 μg mL^−1^ chloramphenicol). Expression cultures were grown at 37°C with 230 rpm shaking to an OD_600_ of 2–2.5 before inducing expression with 0.5 mM IPTG and incubating cultures at 16°C with 230 rpm shaking for 16–20 h. Selenomethionine-labeled protein was produced as previously described by growing expression cultures in modified M9ZB media (2 mM MgSO_4_, 0.4% glucose, 47.8 mM Na_2_HPO_4_, 22 mM KH_2_PO_4_, 18.7 mM NH_4_Cl, 85.6 mM NaCl, 2–50 μM trace metals, 1 μg mL^−1^ thiamine, 100 μg mL^−1^ ampicillin, 34 μg mL^−1^ chloramphenicol)^40^. After expression, 2 L of culture was harvested by centrifugation and resuspended in 120 mL lysis buffer (20 mM HEPES-KOH pH 7.5, 400 mM NaCl, 30 mM imidazole, 10% glycerol, 1 mM DTT). Cells were lysed by sonication then clarified by centrifugation at 25,000 × g for 20 min, and lysate was passed over 8 mL Ni-NTA resin (Qiagen) using gravity chromatography. Resin was washed with 70 mL wash buffer (20 mM HEPES-KOH pH 7.5, 1 M NaCl, 30 mM imidazole, 10% glycerol, 1 mM DTT) and 20 mL lysis buffer, then eluted with lysis buffer containing 300 mM imidazole. Eluate was dialyzed overnight using 14 kDa dialysis tubing in dialysis buffer (20 mM HEPES-KOH pH 7.5, 250 mM KCl, 1 mM TCEP) in the presence of recombinant human-SENP2 to induce SUMO-tag cleavage, before further purification by size-exclusion chromatography using a Superdex 75 16/600 column (Cytiva). Size-exclusion peaks of interest were collected and concentrated to >40 mg mL^−1^, then flash frozen and stored at −80°C.

Ligand-bound Tad1 was produced by first co-expressing Tad1 with BdTIR. BdTIR was cloned into a custom pET vector containing a C-terminal Twin-Strep tag, a chloramphenicol resistance gene, and IPTG-inducible promoter. Plasmids containing cmTad1 or cbTad1 were co-transformed with BdTIR into BL21(DE3) cells, then plated onto MDG agar plates and expressed as described above. Ligand-bound Tad1 was purified as described above with a modified method. After elution from NiNTA resin using 300 mM imidazole, eluate was treated with recombinant human-SENP2 for 1 h to induce SUMO-tag cleavage then immediately purified further by size-exclusion chromatography. Peaks of interest were collected and concentrated to >25 mg mL^−1^, then flash frozen and stored at −80°C. To purify 1′–2′ gcADPR from Tad1, cmTad1 was boiled for 5 min at 95°C, centrifuged for 10 min at 17,000 × g, and filtered through a 10 kDa concentration unit (Amicon). Filtrate was collected, diluted in water, and stored at −20°C.

Crystals of cbTad1 were grown using the hanging drop method using EasyXtal 15-well trays (NeXtal). Sample was prepared by first diluting purified protein to 10 mg mL^−1^ using buffer containing 20 mM HEPES-KOH pH 7.5, 80 mM KCl, and 1 mM TCEP. 2 µL hanging drops were set at a 1:1 ratio of protein to reservoir solution over a well with 400 µL reservoir solution. Each protein was crystallized using the following conditions: 1) Native or selenomethionine-labeled cbTad1 in the apo state: Crystals were grown for 1–2 weeks using reservoir solution containing 0.1 M Tris-HCl pH 7.5 and 40% PEG 200 before being harvested by flash freezing in liquid nitrogen. 2) cbTad1 complexed with 1′–2′ gcADPR: Using cbTad1 purified from cells expressing BdTIR, crystals were grown for 1–5 days using reservoir solution containing 0.1 M MES pH 5.0, 1% PEG-6000, and 300 µM 1′–2′ gcADPR before being cryo-protected with reservoir solution containing 35% ethylene glycol and harvested by flash freezing in liquid nitrogen. X-ray diffraction data were collected at the Advanced Photon Source (beamlines 24-ID-C and 24-ID-E), and data were processed using the SSRL autoxds script (*A. Gonzalez*, Stanford SSRL). Anomalous data for phase determination were collecting using selenomethionine-labeled crystals. Heavy sites were identified, and initial maps were produced using AutoSol in Phenix^41^. Model building was performed in Coot^42^, with refinement in Phenix and statistics were analyzed as presented in Supplementary Table 5^43–45^. Final structures were refined to stereochemistry statistics for Ramachandran plot (favored / allowed), rotamer outliers, and MolProbity score as follows: cbTad1 apo, 98.60% / 1.40%, 0.20%, 1.11; cbTad1–1′–2′ gcADPR, 97.97% / 2.03%, 0.91%, 1.09. See Supplementary Table 5 and ‘Data availability’ for deposited PDB codes. All structure figures were generated with PyMOL 2.3.0.

### Purification and validation of 1′–2′-gcADPR

To purify 1′–2′ gcADPR, concentrated (>1 mM) ligand-bound cmTad1 in storage buffer (20 mM HEPES-KOH pH 7.5, 250 mM KCl, 1 mM TCEP) was denatured by boiling at 95°C for 15 min. Denatured cmTad1 was pelleted at 13,500 × g for 20 min, and supernatant was passed through a 0.5 mL 10 kDa filter (Amicon). 1′–2′ gcADPR concentration was estimated using the extinction coefficient of cADPR and absorbance at 260 nm, which was measured using a Denovix Ds-11+ spectrophotometer.

To validate the signaling capability of 1′–2′ gcADPR isolated from ligand-bound cmTad1, 200 μM ligand-bound cmTad1 was subjected to temperatures of either 25°C or 95°C for 15 min before centrifugation and 10 kDa filtration, and ThsA activity of the resulting filtrate was measured through εNAD fluorescence as described above.

High-performance liquid chromatography (HPLC) was used to further validate the identity of 1′– 2′ gcADPR isolated from ligand-bound cmTad1. 500 μM filtrate was compared to similar amounts of cADPR and ADPR diluted in storage buffer. Individual samples were injected onto a C18 column (Zorbax Bonus-RP 4.6 × 150 mm, 3.5 μm) attached to an Agilent 1200 Infinity Series LC system and eluted isocratically at 40°C with a flow rate of 1 mL min^−1^ with 50 mM Na_2_HPO_4_ pH 6.8 supplemented with 3% acetonitrile. Elution profiles were monitored at an absorbance of 254 nm.

## Data Availability

Data that support the findings of this study are available within the article and its Supplementary Tables and Supplementary files. IMG/MGV accessions, protein sequences and nucleotide sequences appear in Supplementary Tables 2–4. Coordinates and structure factors of cbTad1 apo, and cbTad1–1′–2′ gcADPR have been deposited in the PDB under the accession codes 7UAV and 7UAW.

## Acknowledgements

We thank the Sorek laboratory members for comments on the manuscript and fruitful discussion. We also thank Dr. Yoav Peleg and Dr. Shira Albeck from the Center for Structural Proteomics (ISPC) within the Weizmann Institute of Science for assistance in protein expression, Dr. Morten Danielsen and Daniel Malheiro from MS-Omics for conducting MS experiments, and Dr. Stephen C. Wilson for advice on gcADPR analysis. R.S. was supported, in part, by the European Research Council (grant ERC-AdG GA 101018520), Israel Science Foundation (grant ISF 296/21), the Deutsche Forschungsgemeinschaft (SPP 2330, grant 464312965), the Ernest and Bonnie Beutler Research Program of Excellence in Genomic Medicine, the Minerva Foundation with funding from the Federal German Ministry for Education and Research, and the Knell Family Center for Microbiology. P.J.K. was supported, in part, by the Pew Biomedical Scholars program and The Mathers Foundation. B.R.M. is supported as a Ruth L. Kirschstein NRSA Postdoctoral Fellow NIH F32GM133063 and S.J.H. is supported through a Cancer Research Institute Irvington Postdoctoral Fellowship (CRI3996). X-ray data were collected at the Northeastern Collaborative Access Team beamlines 24-ID-C and 24-ID-E (P30 GM124165), and used a Pilatus detector (S10RR029205), an Eiger detector (S10OD021527), and the Argonne National Laboratory Advanced Photon Source (DE-AC02-06CH11357).

## Supplementary Tables (provided as Excel spreadsheet or Word files)

Table S1. Sequence similarity between alignable regions of SBSphiJ-like phages

Table S2. Candidate genes from SBSphiJ7 that were cloned and tested

Table S3. Tad1 homologs from the IMG database

Table S4. Tad1 homologs from the MGV database

Table S5. Summary of crystallography data collection, phasing, and refinement statistics

## Supplementary Files

File S1. Map of the pSG-thrC-phSpank vector

File S2. Map of the pJG_thrC_dCAS9_gRNA vector

File S3. Map of the pBbA6c_TAD1_His plasmid

File S4. Map of the pBAD_ThsA_ThsB plasmid

File S5. Map of the bdSumo_TAD1 plasmid

## Supplementary Figures

**Figure S1.**
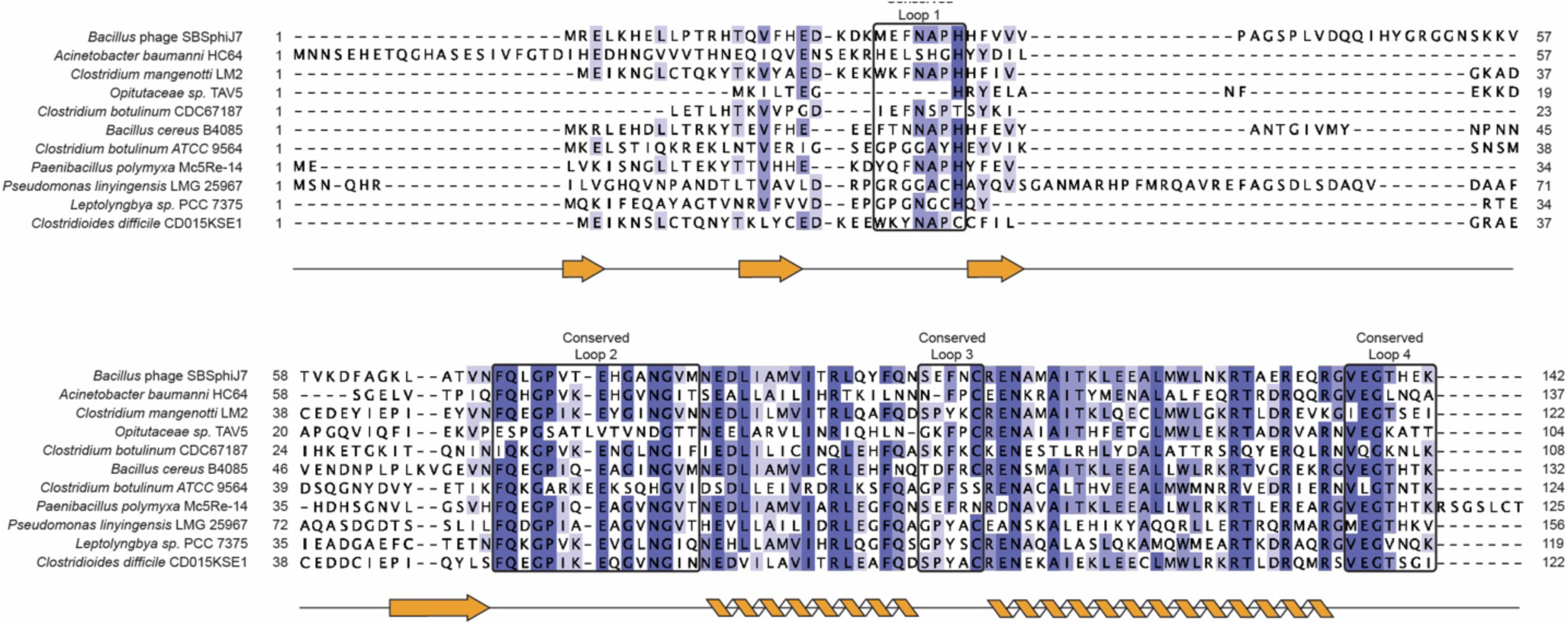
Multiple sequence alignment of the original Tad1 from phage SBSphiJ7, and 10 Tad1 homologs that were verified experimentally as anti Thoeris proteins. The strength of shading indicates degree of residue conservation. The determined Tad1 secondary structure is depicted below and conserved loops involved in ligand-binding are boxed in black.

**Figure S2.**
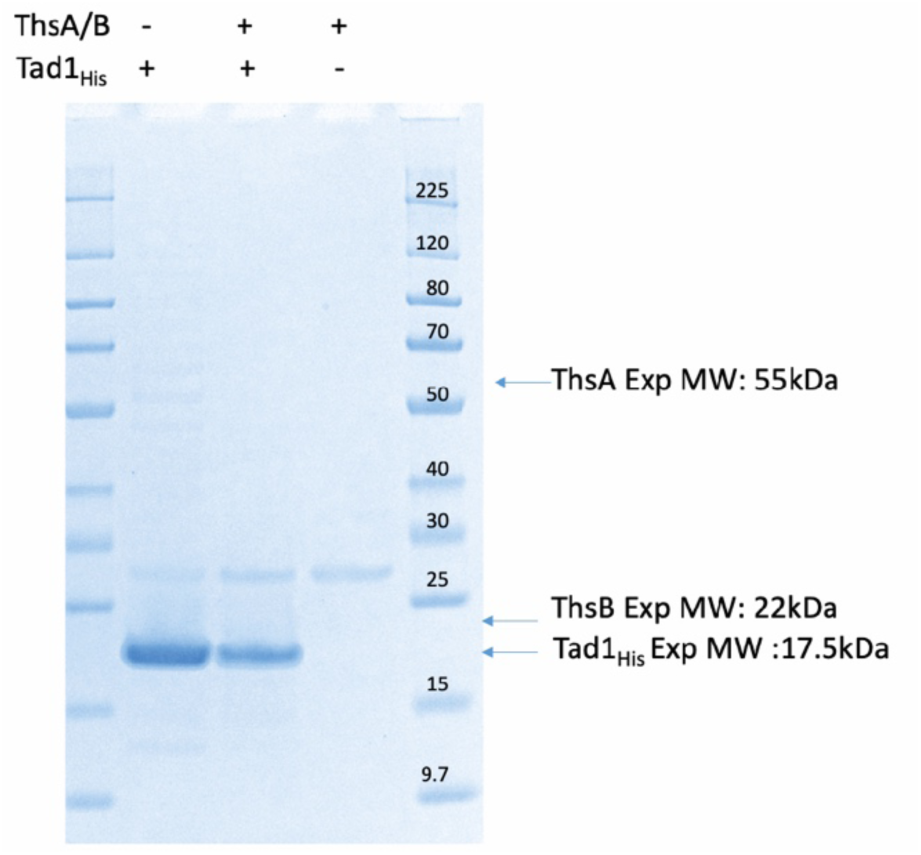
Tad1 does not pull down ThsA or ThsB. Proteins were extracted from *E. coli* cells co-expressing His-tagged Tad1 and Thoeris proteins, and were analyzed on an SDS-PAGE gel. Lane 1: protein size marker. Lane 2: cells expressing His-Tad1 only. Lane 3: cells expressing His-Tad1 and Thoeris proteins ThsA and ThsB. Lane 4: cells expressing Thoeris proteins only. Lane 5: size marker. Expected sizes of His-Tad1, ThsA and ThsB are indicated.

**Figure S3.**
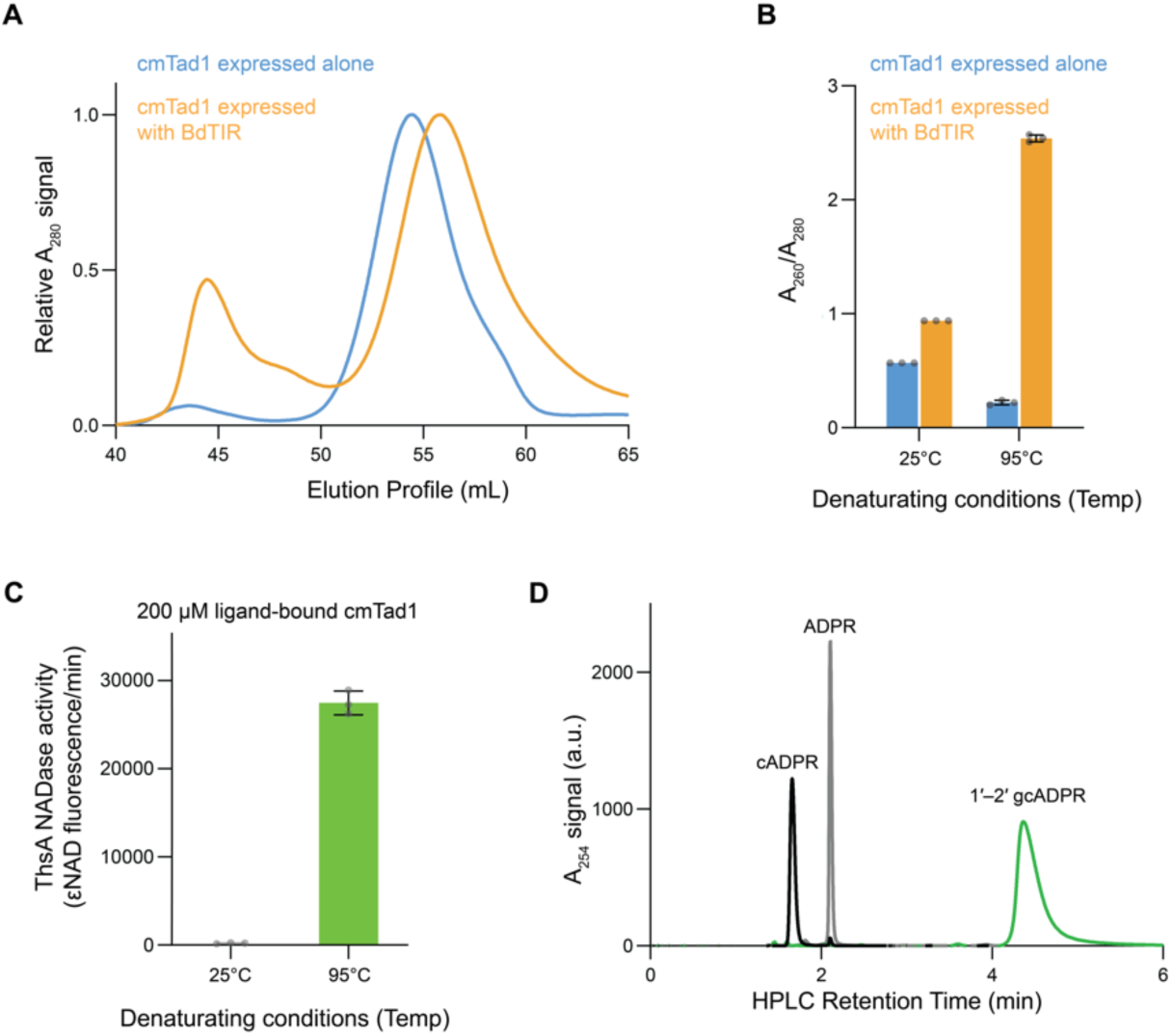
Purification of ligand-bound cmTad1 and 1’–2’ gcADPR. (A) Superdex 75 size-exclusion chromatography of ligand-bound or *apo* state cmTad1 purified from cells expressing BdTIR or only cmTad1, respectively. Ligand-bound cmTad1 shows a ∼0.8 mL right shift compared to cmTad1 in the apo state. (B) A_260_/A_280_ signal ratios of *apo* state and ligand-bound cmTad1 in folding (25°C) or denaturing (95°C) conditions. Folded ligand-bound cmTad1 shows higher A_260_/A_280_ ratio and seems to release a nucleotide species with high A_260_ absorbance upon denaturation. (C) Filtrate derived from 200 μM boiled ligand-bound cmTad1 strongly activates ThsA, while filtrate derived from folded ligand-bound cmTad1 does not. (D) HPLC analysis of filtrate derived from boiled ligand-bound cmTad1 shows a single unique species that migrates distinctly from both cADPR and ADPR.

## Notes

### Competing Interest Statement

The authors have declared no competing interest.

