## Supplementary table 5 for "Viruses inhibit TIR gcADPR signaling to overcome bacterial defense"

**Supplementary Table 5. Summary of crystallography data collection, phasing, and refinement statistics**

|  | cbTAD1  SeMet  **(7UAV)** | cbTAD1–  1′–2′ gcADPR  **(7UAW)** |
| --- | --- | --- |
| **Data collection** |  |  |
| Space group | H 3 2 | H 3 2 |
| Cell dimensions |  |  |
| *a*, *b*, *c* (Å) | 155.95, 155.95, 161.45 | 124.78, 124.78, 113.31 |
| α, β, γ (°) | 90.00, 90.00, 120.00 | 90.00, 90.00, 120.00 |
| Resolution (Å) | 48.67–2.20  (2.27–2.20) | 39.10–1.88  (1.92–1.88) |
| *R*_pim_ | 3.3 (79.3) | 7.0 (57.2) |
| *I* / σ(*I)* | 15.2 (1.5) | 5.3 (1.0) |
| Completeness (%) | 100.0 (100.0) | 99.6 (99.1) |
| Redundancy | 41.5 (41.8) | 5.3 (5.1) |
| **Refinement** |  |  |
| Resolution (Å) | 48.67–2.20 | 39.10–1.88 |
| No. reflections  Total  Unique  Free | 1591383  38341  2000 | 145694  27518  2502 |
| *R*_work_ / *R*_free_ | 20.09 / 24.78 | 16.87 / 19.69 |
| No. atoms |  |  |
| Protein | 4696 (5 copies) | 2006 (2 copies) |
| Ligand / ion | – | 70 (gcADPR) |
| Water | 114 | 306 |
| *B*-factors |  |  |
| Protein | 67.55 | 26.49 |
| Ligand / ion | – | 20.48 |
| Water | 55.51 | 37.05 |
| R.m.s. deviations |  |  |
| Bond lengths (Å) | 0.002 | 0.006 |
| Bond angles (°) | 0.400 | 0.921 |

*All datasets were collected from individual crystals. *Values in parentheses are for the highest resolution shell.
